# *Xanthomonas sontii*, and not *Xanthomonas sacchari*, is the dominant, vertically transmitted core rice seed endophyte

**DOI:** 10.1101/2023.10.19.562881

**Authors:** Rekha Rana, Prabhu B. Patil

## Abstract

Seeds endophytes, particularly the abundant, core and vertically transmitted species, are major areas of focus in the host microbiome studies. Apart from being the first members to colonize, they accompany the plant throughout its development stages and also to the next generation. In a recently published study from china, a *Xanthomonas* species was reported as the keystone species that is core endophyte and vertically transmitted in rice with probiotic properties. However, the species status was wrongly reported as *X. sacchari*. Such report is misleading as *Xanthomonas sacchari* is a well-known and pathogenic species of sugarcane, and the study did not include the two-probiotic non-pathogenic *Xanthomonas* species from rice seeds, *Xanthomonas sontii* and *Xanthomonas indica*, that were discovered and investigated in details by our group, leading to the wrong inference. By including these species, we have correctly established the phylogenetic and taxonomic identity of keystone species as *Xanthomonas sontii*, a non-pathogen with plant protective functions. The course correction will enable researchers to use the correct reference or lab strain of *X. sontii* for further fundamental studies and translational research towards future agriculture.

## Background

The association of microbes with plants, both internally and externally, is a well-established fact. Endophytes are microbes found within the inner parts of plants that typically do not cause any disease symptoms to the host [1]. They play significant roles in plant beneficial functions such as plant growth promotion, inorganic phosphate solubilization, indole-acetic acid (IAA) production, nitrogen fixation, iron sequestering, and plant protection [1-6]. Rice, like any other plant, harbours a large and diverse population of microbes that are associated with different plant parts like leaves, stem, seeds, roots and its micro-environment like rhizosphere [7, 8]. These microbes play an important role in the adaptation and development of plant during its lifetime and also post-harvesting in the form of seed-associated microbes [9, 10]. Seeds are unique niches that allow the microbes to be carried to the next generation and also represent those microbes that traverse from different plant parts into the seeds to be part of the progeny [2, 11].

The advent of the genomic era has enabled us to investigate the identity of the strain/species associated with rice plants at an unprecedented scale and details at the level of taxonomy, ecology and functional capabilities. Our group published the first such study of rice seed microbiome, which led to the discovery of non-pathogenic *Xanthomonas* (NPX) strains which were later described as a novel species, *X. sontii* [12, 13]. Further large-scale isolation of NPX strains from healthy rice seeds and their genomic analysis revealed the presence of another novel but minor species, *Xanthomonas indica* [14, 15]. The genus *Xanthomonas* is primarily known as a pathogen of a wide range of plants infecting in the host and the tissue-specific manner [16]. However, in recent decades, there have been increasing reports of NPX strains from different plants [17]. For example, in rice, only one pathogenic species, *X. oryzae*, has been known for last 100 years [18]. However, in the last one decade, three non-pathogenic species have been reported in rice, two of which are from our lab [13, 14, 19].

Two of the NPX species are from rice seeds, which displayed plant protection functions [15]. Numerous other studies using partial 16S rRNA gene sequencing of metagenomic and cultivation-based approaches have also revealed that *Xanthomonas* is a core vertically transmitted endophyte with plant growth promoting traits such as ACC deaminase activity, IAA production, phosphate and potassium solubilization, siderophore production, as plant growth promoting rhizobacteria (PGPR) and to be involved in early seedling development in submerged conditions that correlates with its reported amylase, cellulase and protease activity [20-22].

In a recently published study, a comprehensive analysis of six different rice varieties from four diverse geographical locations over two generations, revealed that a species of *Xanthomonas* is the core endophyte of rice seeds [22]. Further, the study also provided evidence for the vertical transmission of the core endophytes identified through the study. However, the endophytic *Xanthomonas* strains were wrongly identified as *X. sacchari*, which is a pathogen of sugarcane [23-25]. Apart from our group publications on rice seed microbiome, importantly, the study did not include our earlier publications on novel species of *Xanthomonas* with plant protection properties from rice seed microbiome and associated in-depth studies [13-15]. As the strains from the study [22] are not available in the public domain, researchers interested in understanding this endophyte as keystone species and also exploiting this endophyte will end up working with lab strains or reference strains of *X. sacchari*. Hence, correctly establishing the species status is important for further systematic molecular genetic investigation to understand the mechanistic details leading to its success as a keystone species of rice microbiome.

### Phylogenomic investigation reveals clustering of rice seed endophytic strains with *X. sontii* reference strain

Genome sequences of twenty-seven strains of *Xanthomonas* strains reported by Zhang and co-workers, along with previously reported strains, including type strains of *X*. sontii, *X. sacchari, X. indica* and other species of the genus *Xanthomonas*, were obtained from the NCBI GenBank database [26]. For phylogenetic analysis, genomes were annotated using Prokka v1.14.6 [27] and the GFF files were used to create core gene alignment with Roary v3.12.0 [28]. 407 core genes were used for phylogeny generation with RaxML v8.2.12 with GTRGAMMA model of nucleotide substitution using 1000 bootstrap replicates [29]. The mid-point rooted phylogeny was interactively edited and annotated in iTOL v6 [30]. A core gene-based phylogenetic tree by including the type strains of *Xanthomonas* along with *X. sacchari* strains and non-pathogenic *Xanthomonas* reported from rice seeds, i.e., *X. sontii* and *X. indica* is shown in **Fig. 1**. In the tree, all the twenty-seven endophytic *Xanthomonas* strains formed a monophyletic lineage with *X. sontii* type strain and not with the type strain of *X. sacchari*. Our latest study, reported that the strains of *X. sacchari, X. sontii* and *X. indica* form a sub-clade within the clade I group of Xanthomonads [15]. Hence, to further establish the robustness of the tree, we included all the reported strains of *X. sacchari, X. sontii* and *X. indica*. Both the *X. sontii* and *X. indica* were reported by our group as non-pathogenic *Xanthomonas* species with plant protection properties from rice seeds [15]. As in the dynamics of seed microbiome study, our publications related to non-pathogenic *Xanthomonas* from rice seeds were not cited, and further, the lack of inclusion of the *X. sontii* and *X. indica* type strains led to wrong inference of core-endophytic strains as *X. sacchari*. Furthermore, unlike *X. sontii, X. sacchari* is a well-known pathogen of sugarcane, and its report as a keystone rice seed endophytic species makes no sense. In the phylogenetic tree, it is clear that the strains of *X. sacchari, X. sontii* and *X. indica* make distinct lineages within the sub-clade (**Fig. 1**).

**Fig. 1.**
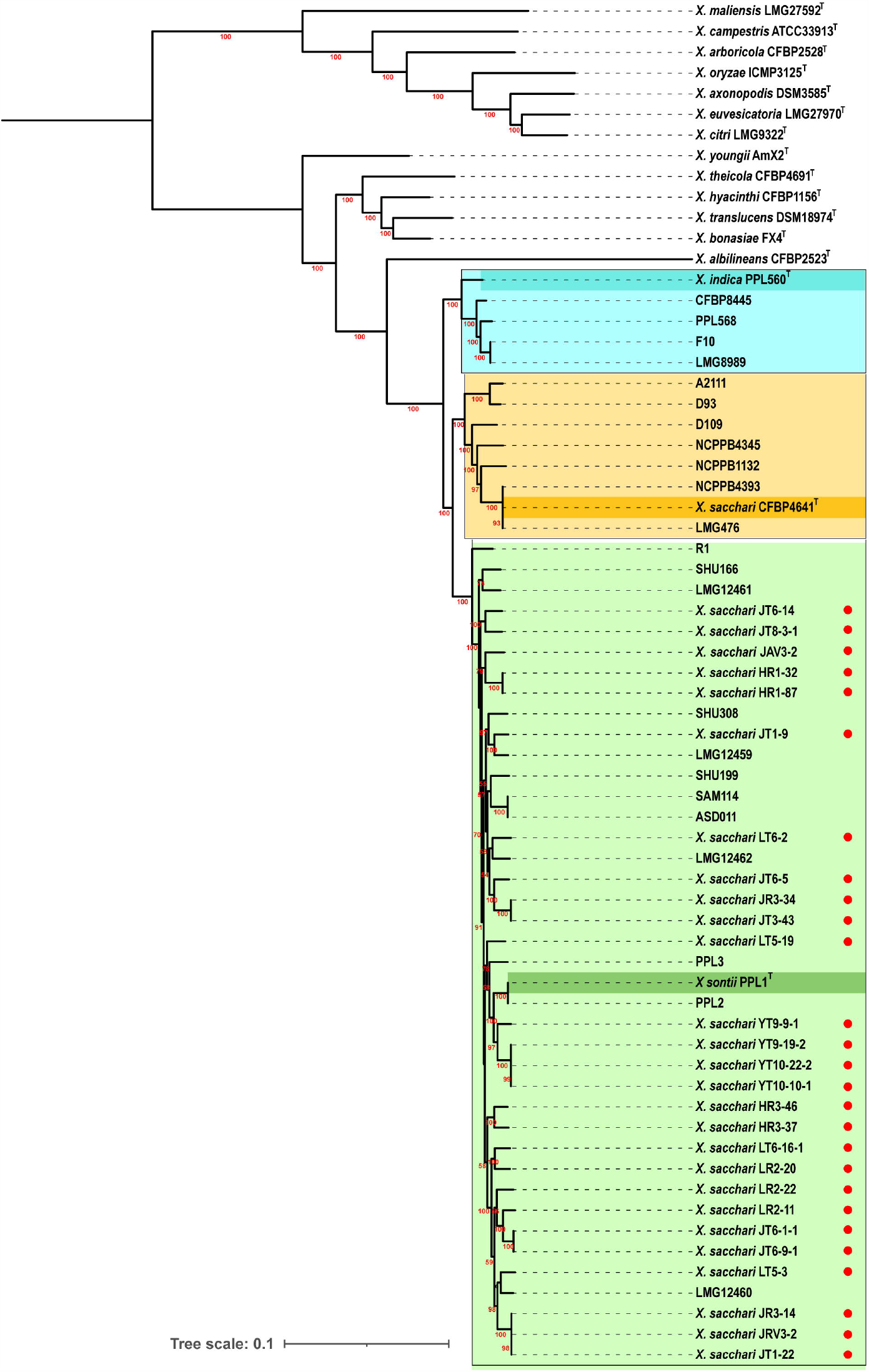
Core gene mid-rooted phylogeny of misclassified *Xanthomonas* isolates with *X. sontii, X. sacchari*, and *X. indica* strains and some other important species of the genus *Xanthomonas*. The red circles in front of the strain names indicate previously misclassified isolates. The three major clades, i.e., *X. indica, X. sontii*, and *X. sacchari*, are labelled in different colours with their type strains highlighted. Bootstrap values are given on the branches. The scale bar denotes the number of nucleotide substitutions per site.

Further, the genome sequences of rice endophytes were submitted to Type (strain) Genome Server (TYGS) for additional validation of phylogenetic clusters [31, 32]. TYGS compares user-submitted genomes against its database of type strains and determines species and subspecies using digital DNA-DNA hybridization (dDDH) values. TYGS determines 10 closely related type strains for each genome submission and constructs a whole genome-based phylogeny using the Genome BLAST Distance Phylogeny (GBDP) approach to calculate intergenomic distances using distance formula d_5_ [33] and the phylogeny is inferred using FastME v2.1.6.1 [34]. The TYGS results shown in **Fig. 2** revealed that all the *Xanthomonas* endophytes are forming a species cluster with *X. sontii* type strain, and *X. sacchari* type strain lies outside in the close association of this species cluster along with *X. indica*, as seen in the core genes based phylogenetic tree **(Fig. 1)**.

**Fig. 2.**
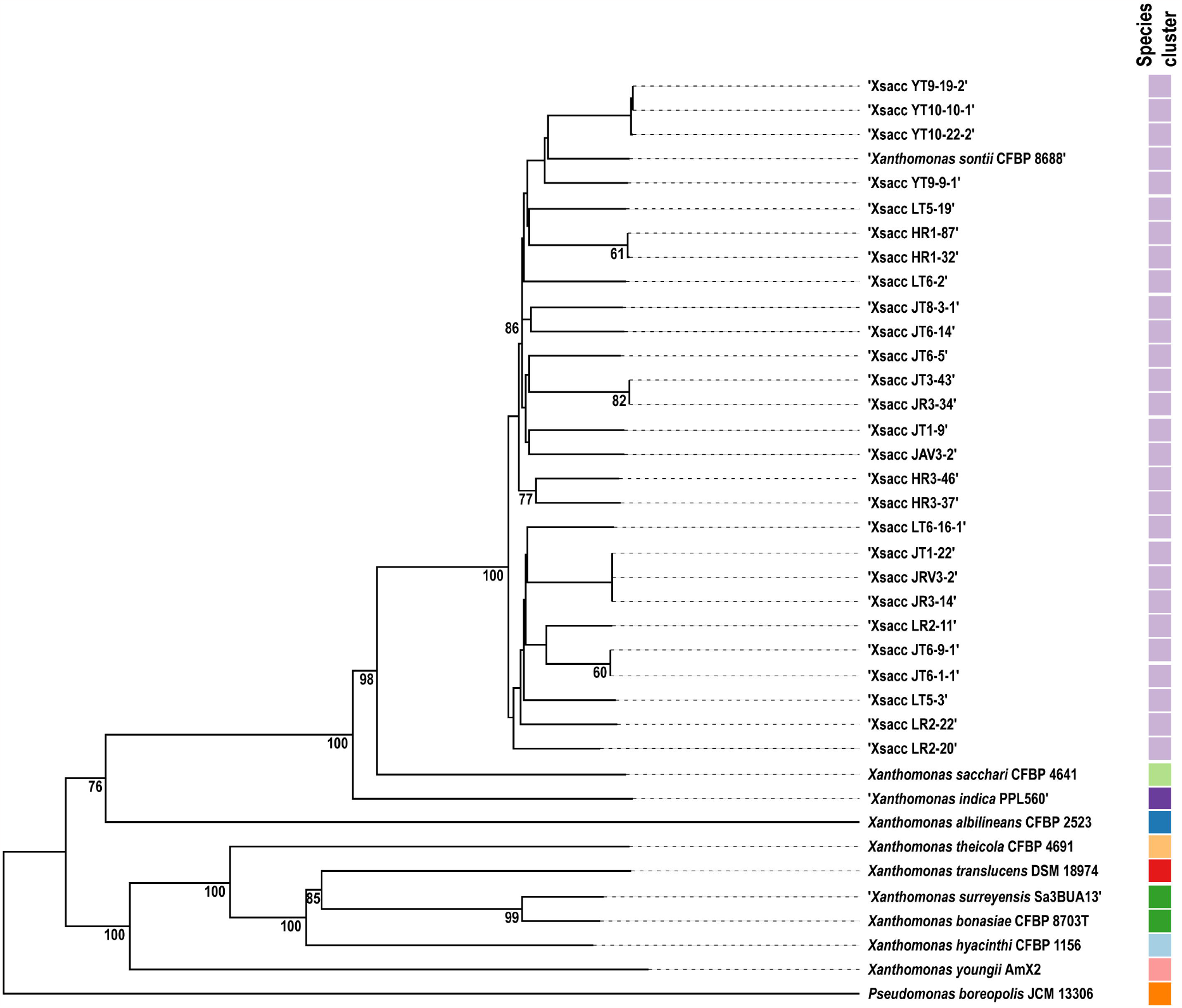
The whole genome-based mid-rooted phylogeny of misclassified *X. sacchari* (Xsacc) isolates constructed by the TYGS web server. The phylogenetic tree was inferred using FastME v2.1.6.1 with GBDP distances using intergenomic distance formula d_5_. The number given above branches denote GBDP pseudo-bootstrap support values for 100 replicates, with an average branch support of 95.5%. Values only above 60% are reported. The coloured bars in front of strain names represent species clusters.

### Taxono-genomics establishes the identity of core endophytic species as *X. sontii*

The Average Nucleotide Identity (ANI) and digital DNA-DNA hybridization (dDDH) values are two important modern criteria for species delineation in this genomic era [35]. To establish the taxonomy, we calculated the ANI values using the OrthoANI algorithm implemented with USEARCH v11.0.667 and dDDH values using formula 2 of Genome to Genome Distance Calculator v3.0 (GGDC) [36, 37]. The ANI values of *Xanthomonas* endophytes were in the range of 97.2 to 98.1% with *X. sontii* type strain, which is much above the cut-off for species delineation and much higher than the ANI values of 94.0 to 94.3% with *X. sacchari* type strain. Additionally, the ANI values of *Xanthomonas* endophytic strains were also calculated against *X. sontii* and *X. sacchari* type strains using JSpeciesWS based on BLAST+ (https://jspecies.ribohost.com/jspeciesws/)[38]. The ANIb values of *Xanthomonas* endophytes with type strains of *X. sontii* and *X. sacchari* were slightly less than orthoANI values (**Table 1**). The ANIb values of *Xanthomonas* endophytes were in the range of 97.0 to 97.8% with *X. sontii* type strain and in the range of 93.4 to 94.0% with *X. sacchari* type strain. Apart from the non-inclusion of *X. sontii* type strain in their study, use of only ANI lead to the wrong inference. Unlike ANI, dDDH is a more robust and discriminatory parameter as it is a distance based genomic related index matrix that does not fragment the sequence during analysis [37]. The dDDH values were 75.4 to 82.2% with *X. sontii* type strain and 54.5 to 55.6% with *X. sacchari* type strain, which is much below the cut-off for species delineation. By including the type strain of *Xanthomonas* species reported from healthy rice seeds, ANI and dDDH data clearly established that all the *Xanthomonas* endophytic strains belong to one species, i.e., *X. sontii* as the ANI and dDDH values are above the cut-off of 95% and 70%, respectively with the type strain (**Table 1**). While dDDH values were not calculated by Zhang and co-workers, the borderline ANI value of 94.7% of *X. sacchari* with *Xanthomonas* endophytic strains might have led to wrong inference. The inclusion of the dDDH value not only clarifies that *X. sontii* is not *X. sacchari* but also clearly establishes its species status as *X. sontii*. The dDDH of the endophytic *Xanthomonas* strains is only around 55% with *X. sacchari* type strains, while it is way above 70% with *X. sontii* type strain (**Table 1**).

**Table 1.** Average Nucleotide Identity (ANI) values calculated using orthoANI and JSspeciesWS (ANIb), and digital DNA-DNA hybridization (dDDH) values of *Xanthomonas* isolates calculated against type strains of *Xanthomonas sontii* and *Xanthomonas sacchari* are given in the table. Type strains of *X. sontii* and *X. sacchari* are highlighted with green and orange, respectively. Misclassified *Xanthomonas* isolates from Zang and co-workers’ study are highlighted with yellow, and previously reported *X. sontii* genomes taken from the NCBI GenBank database are labelled with grey.

| Sr. No. | Strain name | NCBI GenBank Accession number | OrthoANI/ANiB |  | dDDH |  |
| --- | --- | --- | --- | --- | --- | --- |
|  |  |  | *X. sontii* PPL1<sup>T</sup> | *X. sacchari* CFBP4641<sup>T</sup> | *X. sontii* PPL1<sup>T</sup> | *X. sacchari* CFBP4641<sup>T</sup> |
| 1. | HR1-32 | CP099529 | 97.6/97.4 | 94.2/93.6 | 78.3 | 54.9 |
| 2. | HR1-87 | NZ_JANFXE000000000 | 97.6/97.4 | 94.1/93.4 | 78.5 | 55 |
| 3. | HR3-37 | NZ_JANFWT000000000 | 97.5/97.4 | 94.3/94.0 | 77.7 | 55.2 |
| 4. | HR3-46 | CP099531 | 97.4/97.3 | 94.1/93.8 | 77.2 | 54.6 |
| 5. | JAV3-2 | NZ_JANFWN000000000 | 97.4/97.3 | 94.3/93.9 | 77.4 | 55.4 |
| 6. | JR3-14 | CP099534 | 97.3/97.1 | 94.2/93.9 | 75.9 | 54.7 |
| 7. | JR3-34 | NZ_JANFWM000000000 | 97.5/97.3 | 94.1/ 94.0 | 77.4 | 54.5 |
| 8. | JRV3-2 | NZ_JANFWK000000000 | 97.4/97.1 | 94.3/93.9 | 75.9 | 54.7 |
| 9. | JT1-9 | NZ_JANFWU000000000 | 97.4/97.2 | 94.0/93.8 | 77.3 | 54.9 |
| 10. | JT1-22 | NZ_JANFWX000000000 | 97.4/97.2 | 94.2/93.9 | 75.9 | 54.7 |
| 11. | JT3-43 | JANFWL000000000 | 97.4/97.3 | 94.2/94.0 | 77.4 | 55.3 |
| 12. | JT6-1-1 | NZ_JANFWZ000000000 | 97.2/97.0 | 94.1/93.8 | 75.4 | 54.7 |
| 13. | JT6-5 | NZ_JANFXA000000000 | 97.6/97.3 | 94.3/94.0 | 78.3 | 55.1 |
| 14. | JT6-9-1 | NZ_JANFWQ000000000 | 97.2/97.0 | 94.2/93.8 | 75.4 | 54.7 |
| 15. | JT6-14 | NZ_JANFWO00000000 | 97.6/97.4 | 94.2/94.0 | 77.9 | 55.2 |
| 16. | JT8-3-1 | NZ_JANFWP000000000 | 97.5/97.3 | 94.1/93.9 | 77.8 | 55 |
| 17. | LR2-11 | NZ_JANFWV000000000 | 97.2/97.1 | 94.1/93.9 | 75.8 | 55 |
| 18. | LR2-20 | NZ_JANFWW000000000 | 97.3/97.2 | 94.2/94.0 | 76.2 | 55.6 |
| 19. | LR2-22 | NZ_JANFWY000000000 | 97.2/97.1 | 94.1/93.9 | 75.7 | 55 |
| 20. | LT5-3 | NZ_JANFXD000000000 | 97.3/97.1 | 94.2/93.9 | 75.7 | 54.9 |
| 21. | LT5-19 | NZ_JANFXC000000000 | 97.7/97.5 | 94.2/93.7 | 79.1 | 55.1 |
| 22. | LT6-2 | CP099533 | 97.5/97.3 | 94.1/93.7 | 77.6 | 54.9 |
| 23. | LT6-16-1 | CP099532 | 97.4/97.1 | 94.2/93.9 | 75.8 | 54.6 |
| 24. | YT9-9-1 | NZ_JANFXB000000000 | 97.9/97.9 | 94.1/93.7 | 82 | 54.8 |
| 25. | YT9-19-2 | CP099530 | 98.1/97.9 | 94.1/93.6 | 82.2 | 55 |
| 26. | YT10-10-1 | NZ_JANFWR000000000 | 98.0/97.9 | 94.2/93.6 | 82.2 | 55 |
| 27. | YT10-22-2 | NZ_JANFWS000000000 | 98.0/97.8 | 94.1/93.7 | 82.2 | 55 |
| 28. | PPL2 | NQYP000000000 | 99.9/99.9 | 94.2/93.8 | 99.9 | 55.3 |
| 29. | PPL3 | NMPO000000000 | 97.6/97.4 | 94.2/93.9 | 78.6 | 55.2 |
| 30. | LMG12459 | NSGT000000000 | 97.4/97.1 | 94.2/93.8 | 77.1 | 55.1 |
| 31. | LMG12460 | NKUH000000000 | 97.2/97.1 | 94.2/93.9 | 75.5 | 54.7 |
| 32. | LMG12461 | NQYT000000000 | 97.4/97.1 | 94.1/93.8 | 76.3 | 54.9 |
| 33. | LMG12462 | NMPP000000000 | 97.4/97.1 | 94.2/93.7 | 77.6 | 55 |
| 34. | R1 | CP010409 | 96.7/96.0 | 93.9/93.3 | 93.68 | 54.1 |
| 35. | SHU166 | NZ_AQOJ000000000 | 97.5/97.4 | 94.2/93.9 | 78.2 | 55.5 |
| 36. | SHU199 | NZ_AQOI01000000 | 97.5/97.3 | 94.4/93.9 | 78.1 | 55.3 |
| 37. | SHU308 | NZ_AQOK000000000 | 97.5/97.3 | 94.3/93.9 | 77.9 | 55.2 |
| 38. | SAM114 | WJPM000000000 | 97.7/97.4 | 94.2/94.0 | 79.7 | 55.1 |
| 39. | ASD011 | WJPN000000000 | 97.7/97.4 | 94.2/94.0 | 79.7 | 55.1 |

### Frequent misclassification of *X. sontii* isolates as *X. sacchari*

Multiple culture-based and metagenomics studies have identified the genus *Xanthomonas* as one of the most abundant seed endophytes [20, 21, 39]. Similarly, apart from Zang and co-workers, several studies report culturing and highlight the beneficial nature of *Xanthomonas* isolates from rice seeds, as mentioned earlier. However, these strains were misclassified as *X. sacchari* due to the use of limited/limitations of 16S rRNA gene sequence information and also non-inclusion of the type strain of *X. sontii* or non-availability of *X. sontii* type strain when the studies were carried out. Here, we obtained the partial 16S rRNA sequences, from these studies reporting *X. sacchari* as rice endophytes from the NCBI database and assessed them using EZBiocloud [40]. Our analysis revealed that based on the partial 16S rRNA gene sequences the top-hit taxon of these isolates was the *X. sontii* type strain (**Table 2**). Moreover, *Xanthomonas* isolates from rice leaves with plant growth-promoting properties and identified as *X. sacchari* also belong to *X. sontii* based on our EZBiocloud analysis (**Table 2**) [41, 42]. With the new possibilities of complete 16S rRNA sequence metagenomic studies using long-read sequencing technologies and the advent of deep metagenomics and the genomic era can provide much-required resolution and confidence in diversity and culturomics studies. However, the close relationship between *X. sacchari, X. sontii, X. indica* and a recently reported species, *X. hawaiiensis*, that form a clade, it is important to use the right and multiple approaches even when using genomic information to establish the species identity [43]. This is because, in literature, even the genome-based focussed studies have wrongly reported *X. sontii* strain(s) as *X. sacchari* [44, 45]. One such study reported a strain isolated from rice seeds, and the strain was found to have antagonistic activity against rice pathogen *Burkholderia glumae* [44]. Even though isolated from rice seeds, this strain R1 was reported as *X. sacchari*, though it actually belongs to *X. sontii* based on the taxonogenomic analysis [46]. This strain is now being used as a representative of *X. sacchari* in further studies [25, 47, 48]. This instance again highlights the importance of proper taxonomic classification of these misclassified *X. sontii* strains.

**Table 2:**
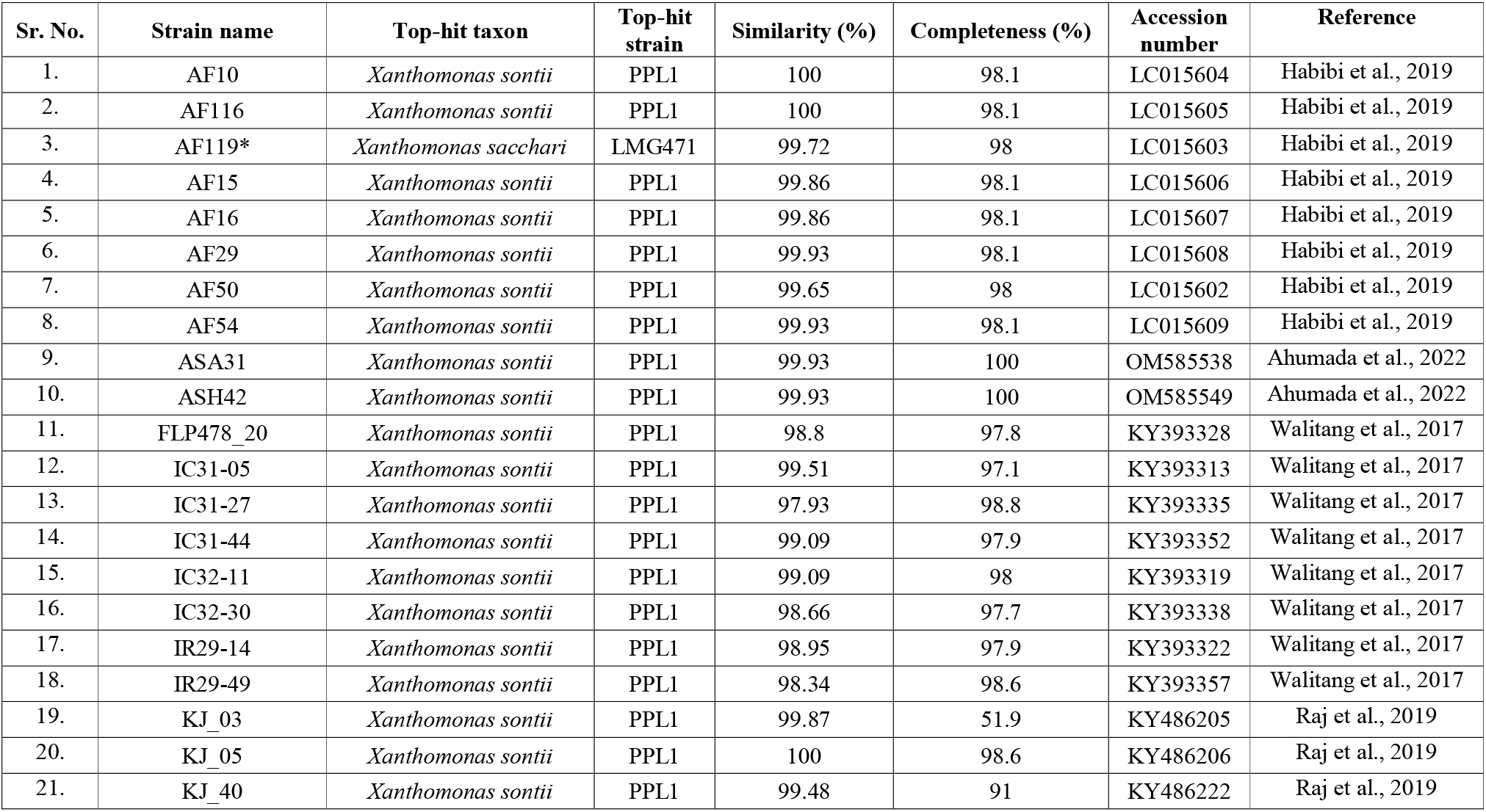
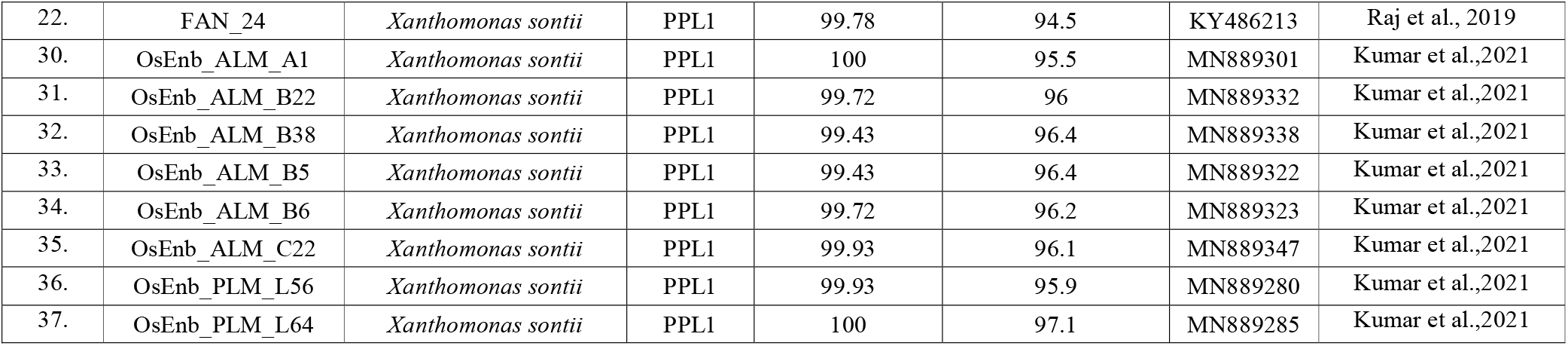
EzBiocloud analysis results of the partial 16S rRNA gene sequences from rice plant isolates wrongly reported as *Xanthomonas sacchari* in previous studies against the EzBiocloud 16S database. The table shows the top-hit taxon and top-hit strains for each 16S rRNA sequence, along with its completeness and similarity to the top-hit strains. The NCBI accession numbers for each sequence are given with references. *Strain which have *X. sacchari* type strain as top-hit taxon.

There are many instances of classifying *X. sontii* strains as *X. sacchari* strains based on ANI values and not considering the dDDH [22]. Even in the follow-up in-depth publication, they have not corrected the mistake and to referred the endophytic strain as *X. sacchari* instead of *X. sontii*. This follow-up study is particularly important as they report that the effect of early inoculation of the endophyte as a keystone species on alterations in microbial composition, diversity, network structure and co-enrichment of bacterial genera well documented for plant beneficial functions [49]. These taxonomic mistakes would hinder further exploitation of *X. sontii* as probiotics apart from investigating the genetic basis of its success as a core endophyte. If these non-pathogens keep getting identified as *X. sacchari*, regulatory procedures might mistake them for harmful pathogens, which can hamper global agricultural export-import of rice seeds. Considering the robustness and power of dDDH as the genomic index in species delineation, new servers have been developed to assist the researchers in typing and classification of strains. At the genomic level, *X. sontii* and *X. sacchari* are closest relatives, and large-scale comparative genomic studies could help find some genetic markers for their proper identification and also to find genomic features that are making *X. sacchari* a successful pathogen of sugarcane when its genome bears such resemblance to a non-pathogenic core rice seed endophyte.

## Conclusion

The study by Zhang and co-workers’ is a pioneering in the field of rice microbiome in general and rice seed microbiome in particular. It is also a major addition to the studies in the emerging field of non-pathogenic *Xanthomonas* with probiotic properties. Hence, the report of *X. sacchari* instead of *X. sontii* as a predominant core-vertically transmitted seed endophyte has broad implications. Even in a follow-up but major study to understand how early inoculation of a core-rice seed endophyte alters the microbiome, the authors have not corrected *X. sacchari* as *X. sontii*. The plant pathogenic *Xanthomonas* species, i.e., *X. oryzae*, has been a model bacterium for host-pathogen interaction, leading to several fundamental discoveries in the field [18]. In this context, the identification of a *Xanthomonas* species as an abundant, core, vertically transmitted seed endophyte and our establishment of its species status as *X. sontii*, a non-pathogenic *Xanthomonas* species with plant protective properties [15] will allow to layout a path for molecular studies of *X. sontii* as a model bacterium for rice (host)-*X. sontii* (microbe) interaction. There is a possibility of such mistakes at the taxonomic level in studies towards the identification of core-commensal or endophytic studies in such studies from other plants and similar studies in human microbiome. Hence, systematic and comprehensive taxonogenomic studies are critical in both metagenomic and culturomics-based microbiome focused studies on probiotic strains.

## List of abbreviations

NPX: Non-Pathogenic *Xanthomonas*
ANI: Average Nucleotide Identity
dDDH: digital DNA-DNA hybridization

## Declarations

### Ethics approval and consent to participate

Not applicable

### Consent for publication

Not applicable

### Availability of data and materials

Not applicable

### Authors’ contribution

RR designed the study, carried out genomic and taxonomic analysis, and drafted the manuscript. PBP participated and coordinated in design of study, applied for funding, and finalised the manuscript.

## Acknowledgements

Not applicable

